# Contribution of local recombination and AT-biased mutations to differentiated region formation in *Apis cerana*

**DOI:** 10.1101/2024.09.02.610918

**Authors:** Fushi Ke, Liette Vasseur

**Affiliations:** Guangzhou, China; Department of Biological Sciences, Brock University, St. Catharines, ON, Canada

**Keywords:** Eastern honeybee, low GC, mutation rate, mutation load, recombination

## Abstract

Genome architecture can interact with evolutionary processes and be involved in the formation of differentiated regions potentially containing adaptation and speciation loci. Regions of low GC content can be linked to low mutation and recombination rates. These effects on the formation of heterozygous differentiation landscapes have not been investigated. Here, we explored mutation accumulation and group divergence along a speciation continuum using 499 genomes of *Apis cerana*, with a widely distributed Central group diverged with its peripheral groups at both population genetic and phylogenetic timescales. We found that the differentiated regions had generally lower recombination and GC content than the rest of the genome, with lower-than-average divergence (*d*_xy_) initially to higher-than-average ones at deeper timescale. Most rare alleles (∼80%) are AT-biased that derived from GC, resulting in lower AT-biased mutations in low-GC regions. Multiple regression analysis further shows that reduced mutation and recombination rates are associated with decreased diversity, particularly in regions with low GC content. Moreover, in all *A. cerana* groups, higher mutation load and less efficient selection in low-GC regions compared with high-GC regions suggest that restricted recombination is important in mutation accumulation (e.g., AT-biased) and group divergence. This pattern explains the increased *d*_xy_ in low-GC regions over evolutionary time. Finally, low-GC regions possess higher proportion of group-specific polymorphisms, which reconciliate discordance between mitochondrial and nuclear phylogenies in *A. cerana*. Our results highlight the contribution of genome architectures to the formation of differentiation landscapes along divergent groups, emphasizing caution regarding loci identified solely from differentiated regions.

**Significance Statement:** This study shows the low nucleotide diversity in differentiated regions could be simply attributed to genome architectures (e.g., gBGC, local recombination and mutation rates). Further studies should consider the effects of genome architecture on understanding the formation of heterozygous differentiation landscape along diverging groups and identification of adaptive loci.

## Introduction

Genomic islands are genomic regions with reduced nucleotide diversity (π) and elevated genetic differentiation (*F*_ST_) that may potentially be involved in adaptation and reproductive isolation (Turner et al. 2005). Understanding the formation of these divergent peaks is imperative in defining how new species arise (Han et al. 2017; Wang et al. 2019; Wolf and Ellegren 2017). While modern sequencing technologies have facilitated our understanding of heterogeneous genomic landscapes in many species (Han et al. 2017; Ravinet et al. 2017; Vijay et al. 2016; Wang et al. 2016, 2019; Wolf and Ellegren 2017), the causes of genomic islands against the troughs of differentiation across the genome remains controversial (Cruickshank and Hahn 2014). Increasing evidence shows that genomic islands may emerge by processes unrelated to adaptation and speciation (Cruickshank and Hahn 2014; Guerrero and Hahn 2017; Ravinet et al. 2017; Wolf and Ellegren 2017). Authors (Burri 2017; Cruickshank and Hahn 2014; Guerrero and Hahn 2017; Han et al. 2017; Ravinet et al. 2017; Vijay et al. 2016; Wang et al. 2016, 2019; Wolf, Ellegren 2017) have proposed multiple evolutionary models to explain the formation of genomic islands in evolving groups. Recurrent selection can increase *F*_ST_ while reducing nucleotide divergence (*d*_xy_) in genomic islands (Charlesworth et al. 1993; Irwin et al. 2016; Ravinet et al. 2017; Burri 2017). Selection in allopatry can generate regions with elevated genetic differentiation (with reduced nucleotide diversity), while not affecting their nucleotide divergence (Cruickshank & Hahn, 2014; Irwin et al., 2018). On the other hand, divergent selection coupling with gene flow may lead to the formation of genomic islands with both elevated *F*_ST_ and *d*_xy_ (Feder et al. 2012; Wolf and Ellegren 2017). Genomic islands with high *F*_ST_ and *d*_xy_ can also arise from ancient polymorphism, especially if gene flow is absent (Guerrero and Hahn 2017; Noor and Bennett 2009). In addition, genome architectures, such as gene density, recombination, and mutation rate, can all contribute to the formation of heterozygous differentiation landscape along the speciation continuum (Burri 2021; Castellano et al. 2020; Leroy et al. 2021; Ravinet et al. 2017).

Unlike other genome architectures, few have investigated the contribution of GC content (proportion of guanine (G) and cytosine (C) nucleotides) on mutation accumulation and genetic differentiation along diverging groups. While GC content may not be a direct driver of differentiation or divergence, positive correlation between GC content and regional recombination rate has been reported from yeast to human (e.g. Meunier et al. 2004; Pessia et al. 2012; Kiktev et al. 2018). Regions with high-GC content are more susceptible to double-strand DNA breaks (i.e., recombination) than low-GC ones (Petes 2001; Kiktev et al. 2018; Bouwman et al., 2023). On the contrary, recombination can drive the fixation of G and C alleles through GC-biased gene conversion (gBGC) (Charlesworth et al., 2020; Fullerton et al. 2001; Meunier et al. 2004; Pessia et al. 2012).

Through generation of AT mutations (Hershberg and Petrov, 2010; Lynch 2007), GC content may increase the local mutation rate (Fryxell and Moon 2005; Kiktev et al. 2018, Long et al., 2018). Rare, derived alleles can be used as a proxy of mutation rate since natural selection has yet to act on them in the short evolutionary time (Coventry et al., 2010; Keinan and Clark, 2012). In *Apis mellifera* genome, for example, elevated C/G to A/T mutations account for over 80% of rare, derived alleles (derived allele frequency < 1%) (Wallberg et al., 2015). Regional high GC would therefore contribute to elevated local mutation rate. Simulation results in *Drosophila melanogaster* show that local mutation rate is a crucial factor interacting with other forces, such as recombination, to shape nucleotide diversity and thus differentiation across the genome (Barroso and Dutheil 2023). Generation of more AT alleles can be further counteracted by gBGC that leads to fixation of more GC alleles (e.g., Wallberg et al., 2015). High recombination (e.g., in high GC regions) can lead to more efficient purified selection by removing deleterious mutations (Hill and Robertson, 1966; MacPherson et al., 2021). These effects may be prominent in genomes with high recombination, such as the honeybee genome, which has an average rate of 19 cM/Mb (centimorgans per megabase pair)—approximately 10 times higher than that of most eukaryotes (Beye et al., 2006). Nevertheless, the interplay between three processes—the generation of AT-biased mutations, the fixation of GC alleles via gBGC, and the removal of deleterious mutations—varies significantly across the genome due to the heterozygous genome architectures (e.g., local GC content and recombination). This interplay may contribute to GC dynamics across the genome and formation of heterozygous differentiation landscape along the speciation continuum.

*Apis cerana* is a widespread cavity-nesting honeybee group (Ji et al. 2020; Qiu et al. 2023; Radloff et al. 2010; Su et al. 2023). Its distribution in Asia ranges from Pakistan to Japan and extends from the adjacent regions of Siberia in the North to its southern distribution in Indonesia, from where the bee swarms invaded Oceania (Koetz et al. 2013). There are four putative species in this group based on the inference of genome-wide single nucleotide polymorphisms (SNPs) (Su et al. 2023): *A. cerana* (Asian Mainland and Indian black, Lineage M), *A. cerana* (Sundaland, Lineage S), *A. cerana* (Indian yellow, Lineage I), and *A. cerana* (Philippines, Lineage P). The mainland lineage (Lineage M) is the most widely distributed species and formed several peripheral groups/populations with varied levels of divergence at the surrounding of its Central population (Ji et al. 2020; Qiu et al. 2023). The Central population of Lineage M admixes with the Lineage M peripheral populations and introgresses with the other *A. cerana* species in the South (Lineage S) (Ji et al. 2023), forming multiple admixed group pairs diverged at both population genetic and phylogenetic timescales.

Here, we used resequencing genomes of 499 *A. cerana* and six *A. mellifera* individuals (outgroup), with groups diverged over evolutionary time, to investigate the contribution (direct or indirect) of GC content to genome-wide differentiation landscape and mutation accumulation over time. Across the genome, GC content (negative), recombination (positive) and mutation rates (positive) are the best variables explaining genome-wide variation of π. When dividing the genome into regions with different GC content, we found that low GC regions were always with lower π and higher *F*_ST_ (i.e., genomic islands) than the rest of the genome. These regions had few AT-biased mutations (the most dominant mutation type, from ancestral G/C to derived A/T alleles) due to lower GC content, which suggested lower mutation rate. In addition, lower recombination rates in these regions are associated with reduced effects of genetic reshuffling on π. Groups at earlier stages of divergence (i.e., population genetic timescale) showed low *d*_xy_ but high *F*_ST_ in low GC regions, which meant even a small number of different mutations between groups could cause high *F*_ST_ if π was low. The groups that were more diverged exhibited elevated levels of *d*_xy_ due to the build-up of group-specific mutations (such as fixed AT alleles), leading to *d*_xy_ values at the phylogenetic timescale (between species) to be greater than the genome-wide average. Higher mutation loads and less efficient selection in low GC regions of all investigated *A. cerana* groups supported the important role of restricted local recombination in variant accumulation and group divergence in these regions. Conflict between mitochondrial phylogeny (Qiu et al. 2023) and its nuclear counterpart (Ji et al. 2020) in *A. cerana* Lineage M could be due to the above-mentioned admixture between adjacent groups. They were reconciled with polymorphisms in low GC regions, where more group-specific variants accumulated. These group-specific variants were easily found in regions with elevated differentiation (i.e., *F*_ST_) and divergence (i.e., *d*_xy_, Central *vs*. Lineage S) but possibly not related to adaption and speciation processes.

## Results

### Genetic structure and divergence of Apis cerana

In ADMIXTURE analysis, the optimal K value (i.e., genetic clusters) was 14 when investigating the genomic polymorphisms of 499 *A. cerana* individuals (Figure 1, Supplementary Table 1). *A. cerana* Sundaland individuals (samples from Malaysia and Singapore, Lineage S) first separated from the mainland species (Lineage M) due to long-term divergence (Su et al. 2023). Island groups (e.g. Taiwan, Hainan and Japan) and populations that colonized extreme habitats (e.g. high elevation: Ind_Pak, Bomi, Aba and Qinghai; or high latitude: Northeast) gradually separated. These groups had clear genetic ancestry and geographical distribution (Figure 1). Environmentally differentiated habitats also contributed to the formation of two clusters in the Central group, and four clusters in the Japan group (Figure 1). These clusters displayed intensive gene flow with adjacent ones and were difficult to distinguish from each other (Figure 1). Based on geography, we grouped the two clusters from Central and the four from Japan into one group. In total, we identified ten groups based on genomic clustering and geography. We performed the following analysis with this grouping scheme. It is worth noting that the grouping scheme is not associated with subspecies identification in the eastern honeybee (e.g., Qiu et al., 2022), but for the convenience of data analysis and illustration. It is important to note that the investigated genomic variation and differentiation pattern, as well as divergence time estimation (see below) when combining two Central clusters as one group (Central group) was consistent with the pattern using either one or the other cluster. Clustering of bioclimatic variables could not clearly separate these groups with some overlapping, suggesting a combination of phylogeny, genetic drift, and natural selection in the formation of these groups. The Central group (of Lineage M) was widely distributed and showed varied extent of genetic admixture with peripheral groups of Lineage M and Lineage S (Figure 1). It was thus used as an invariable group for divergence estimation in a time series (see below).

**Table 1.** Regression analysis of nucleotide diversity on genomic architectures.

| Regions | R<sup>2</sup> | Adj.R<sup>2</sup> | GC content | recombination | Gene density | DAF_001 |
| --- | --- | --- | --- | --- | --- | --- |
| Genome wide | 0.6856 | 0.685 | -2.341*** | 1.648*** | -0.07695* | 2.174*** |
| Low-GC | 0.7715 | 0.7706 | -0.2402*** | 0.5548*** | -0.0995*** | 0.5360*** |
| High-GC | 0.5543 | 0.5526 | -0.9372*** | 0.0182(NA) | -0.0928*** | 0.5788*** |
| region1 | 0.8465 | 0.8443 | -0.0915* | 0.4214*** | -0.1700*** | 0.6128*** |
| region2 | 0.644 | 0.6413 | -0.2717*** | 0.3638*** | -0.2496*** | 0.4904*** |
| region3 | 0.4621 | 0.4591 | -0.4429*** | -0.1075** | -0.1495*** | 0.6985*** |
| region4 | 0.4301 | 0.425 | -0.5802*** | 0.1550*** | -0.0091(NA) | 0.4222*** |
| region5 | 0.438 | 0.4054 | -0.4521*** | 0.1446(NA) | 0.1490(NA) | 0.5408*** |

In addition to low-GC and high-GC regions, genomic windows (100 kb) were divided into five regions (region 1 -5,
from low to high GC) based on equal-width intervals of GC content. Nucleotide diversity ( $\pi$ ) and
recombination (i.e., population-scaled recombination rate) are PC1s extracted from six *A. cerana* Lineage M
groups, respectively. DAF\_001, as a proxy of mutation rate, is number of SNPs with rare and derived alleles
(d<sub>af</sub> < 0.01) in Lineage M. $DAF_{001} \propto \mu$ and $recombination = 3 * N_e * c$ ( $N_e$ , effective population size; $\mu$ ,
mutation rate; $c$ , recombination rate). NA, \*, \*\*, \*\*\* indicate $p > 0.05$ , $p < 0.05$ , $p < 0.01$ and $p < 0.001$ ,
respectively.

**Figure 1.**
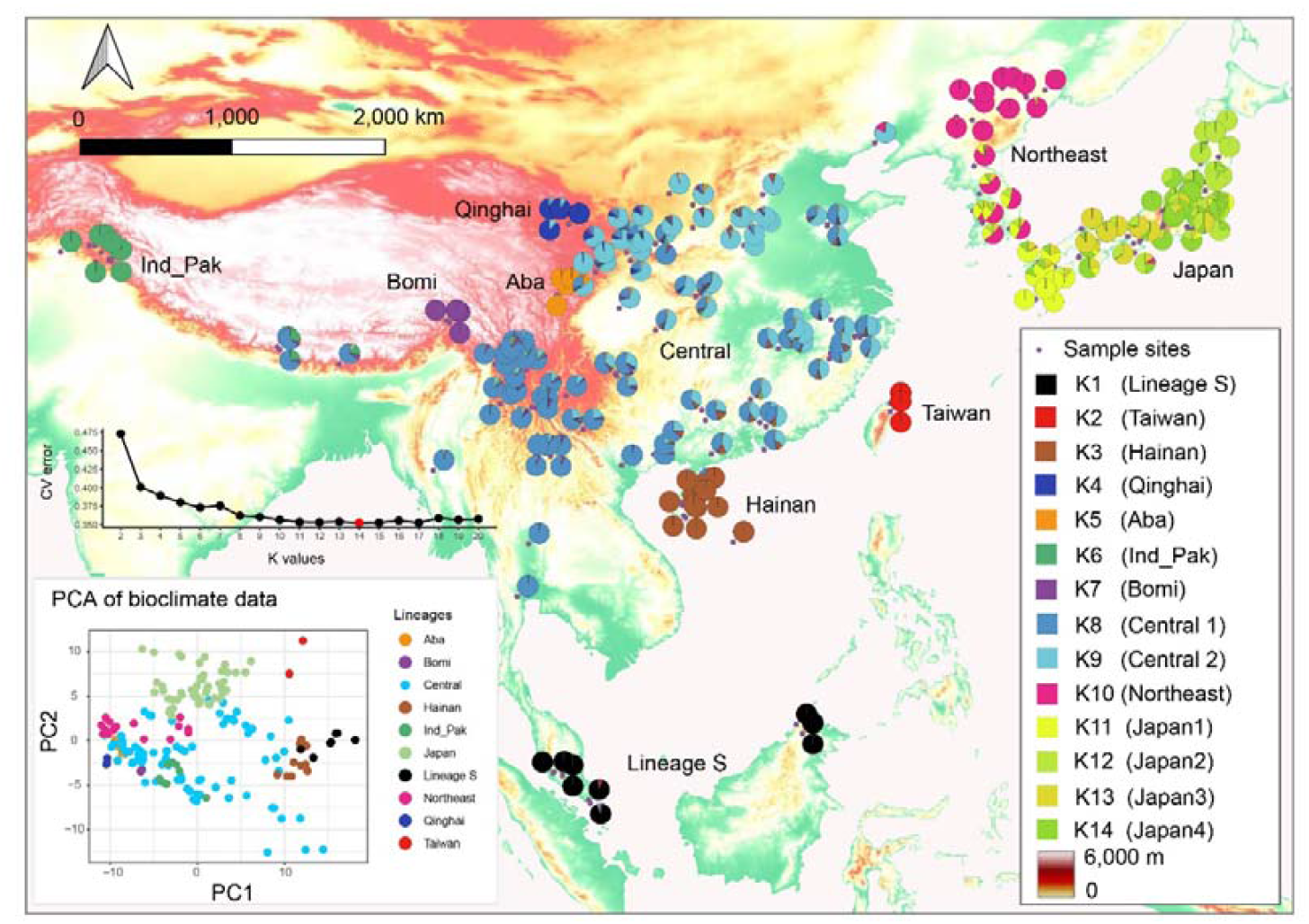
Genetic structure and bioclimate variable clustering of *Apis cerana* Lineage M (mainland lineage). Additional samples from Malaysia and Singapore (i.e., Sundaland lineage (Lineage S); Su et al. 2023) were included. The optimal K=14 was selected based on cross-validation error in ADMIXTURE. Two and four genetic clusters were identified from Central and Japan groups, respectively. The 144 monthly bioclimate variables can be found in Supplementary Table 1.

Estimated divergence times of group pairs within *A. cerana* Lineage M were 502 thousand years (Kyr) in Central *vs*. Japan, 521 Kyr in Central *vs*. Qinghai, 653 Kyr in Central *vs*. Bomi, 700 Kyr in Central *vs*. Hainan, 972 Kyr in Central *vs*. Taiwan (Supplementary Table 2). The divergence was consistent for different group pairs whether using individuals from Southern or Northern part of Central group except for the Central with Japan pair, where individuals from Northern part of Central group showed shallower divergence (Supplementary Table 2). The gradually divergent pattern of five groups with Central group was consistent with the topologies based on nuclear (see below) and mitochondrial polymorphisms (Qiu et al., 2022), thus supporting the robustness of divergence time estimation. These time nodes within population genetic timescale (t1: Central *vs*. Japan; t2: Central *vs*. Qinghai; t3: Central *vs*. Bomi; t4: Central *vs*. Hainan; t5: Central *vs*. Taiwan) and two additional ones from the phylogenetic timescale (t6: Central *vs*. Lineage S; t7: Central *vs*. A. mellifera) were chosen (Supplementary Table 2) and used in a time series to investigate changes of genomic differentiation landscape and characterize genomic polymorphisms.

### Landscape of genomic variation and differentiation of diverging group pairs

By investigating genomic variation (nucleotide diversity, π) and differentiation (genetic differentiation, *F*_ST_ and nucleotide divergence, *d*_xy_) landscape over time (Figure 2A, Supplementary Figure 1-3), we found a distinct pattern between high GC and low GC regions, not just between centromere and non-centromeric regions. Taking *d*_xy_ in low GC regions, for example, divergence increased from t1 to t7, with *d*_xy_ values in t6 and t7 higher than the genome-wide average (Figure 2A, Supplementary Figure 1). In high GC regions, however, divergences over time were indistinguishable among the different group pairs when compared with low GC regions in the genome-wide context (Figure 2A). We further compared nucleotide divergence in low GC and high GC regions over time and found significantly lower *d*_xy_ values in low GC regions from t1 to t4 than in high GC regions (Figure 2B, Mann-Whitney U test, *p* < 0.001). In t5, t6 and t7, however, *d*_xy_ values in low GC regions were significantly higher than in high GC regions (Figure 2B, Mann– Whitney U test, p < 0.001).

**Figure 2.**
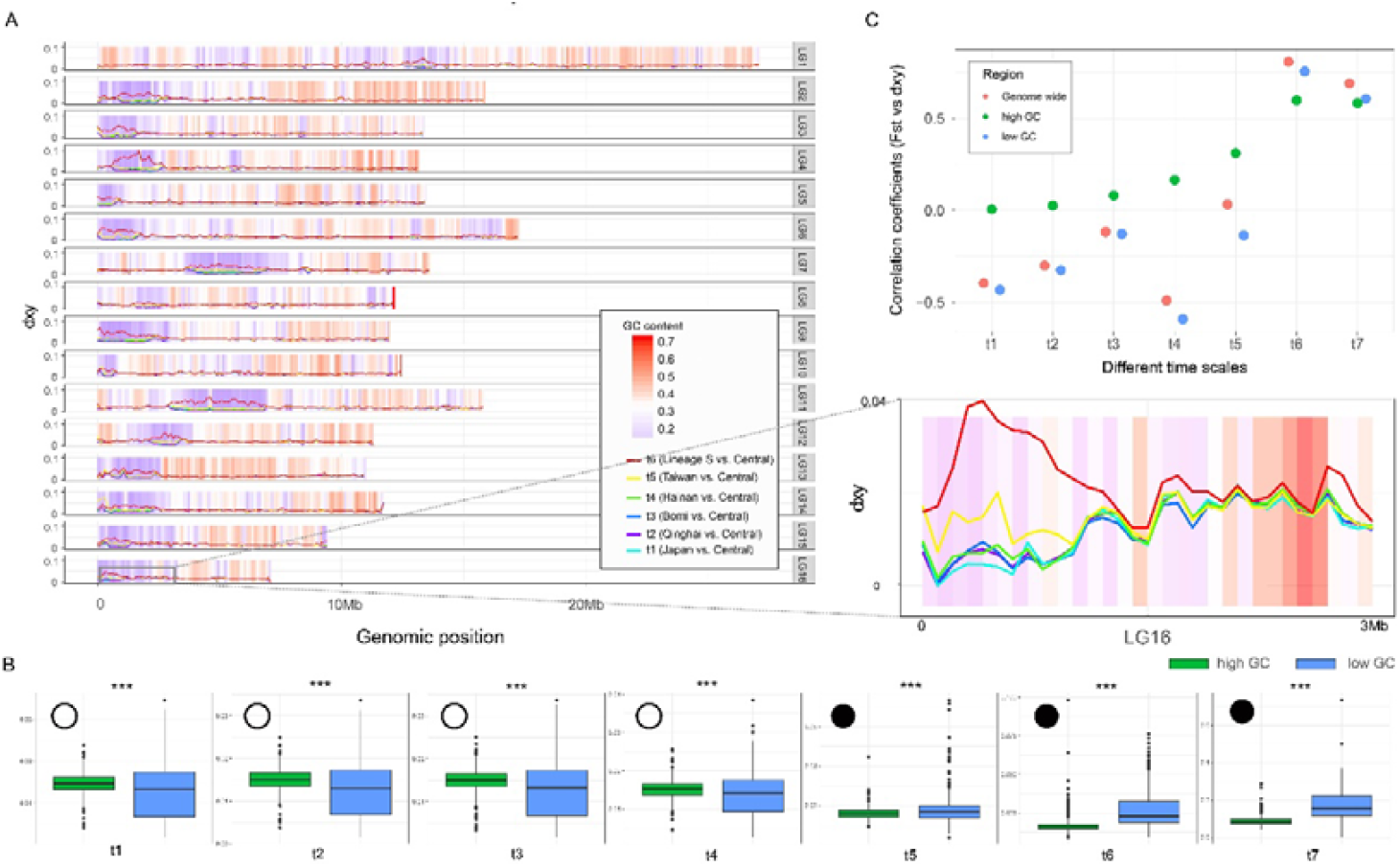
Characterization of heterozygous divergence (*d*_xy_) landscape in high GC and low GC regions. **(A)** Landscape of *d*_xy_ over time (t1 to t6) varied along GC gradient. Landscape of *d*_xy_ from t1 to t7 in Supplementary Figure 1; **(B)** Patterns of *d*_xy_ in low GC and high GC regions shift over divergence time (i.e., progressing from t1 through t5 (population genetic timescale) to t6 and t7 (phylogenetic timescale); Wilcoxon rank sum test was conducted with values between low GC and high GC regions. Empty circles represent significantly lower values in low GC than high GC regions, while black circles indicate low GC regions with higher values. **(C)** Correlation coefficients of *d*_xy_ and genetic differentiation (*F*_ST_) varied over divergence time in different genomic regions.

Correlations between *d*_xy_ and *F*_ST_ (Figure 2C) also switched over time in low GC regions, with a negative correlation from t1 (*r* = -0.43) to t5 (*r* = -0.14), then positive in t6 (*r* = 0.75) and t7 (*r* = 0.61). This pattern was not found in high GC regions (Figure 2C, all with positive r values). Correlation between *d*_xy_ and recombination rate (rho) in low GC shifted pattern while not in high GC regions (Supplementary Figure 4A), with positive r values from t1 (0.41) to t5 (0.26) and negative *r* values in t6 (-0.64) and t7 (-0.48), and a similar trend found in correlations between *d*_xy_ and GC content (Supplementary Figure 4B). In addition, we found both nucleotide diversity (π) and mutation load showed positive correlations with GC content in low GC regions, but negative correlations in high GC regions (Figure 3; Supplementary Figure 4). Consistent negative correlations between nucleotide diversity (π) and gene density were detected across the genome, as well as in high GC and low GC regions (Supplementary Figure 4), supporting widespread linked selection in the functional elements.

**Figure 3.**
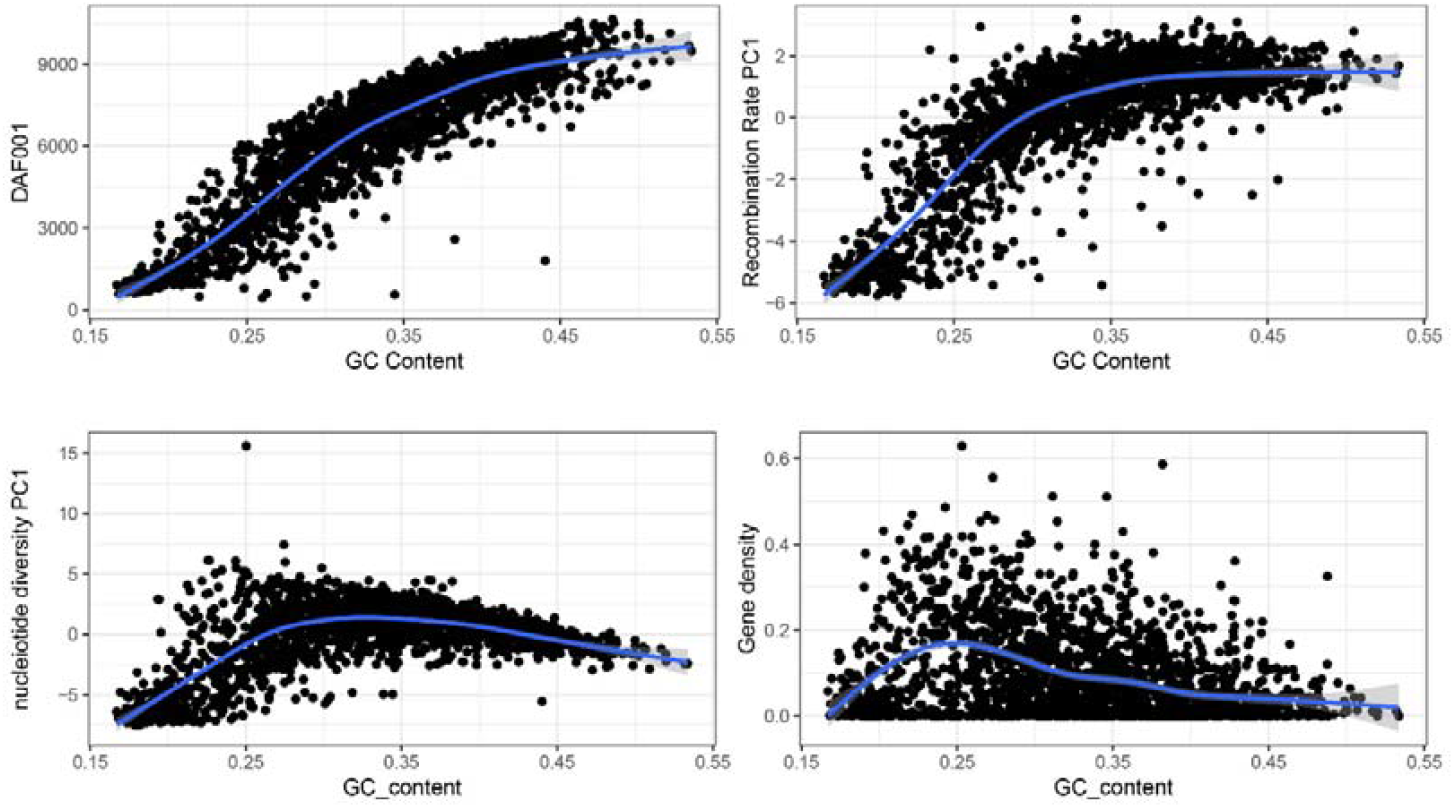
Correlation of GC content with different population genetic parameters (DAF_001, nucleotide diversity PC1, recombination rate PC1 and gene density) of *A. cerana* Lineage M. DAF_001 represents derived allele frequency (DAF) ≤ 0.01. Nucleotide diversity PC1 (explained 86.02% variance) and recombination rate PC1 (explained 74.91% variance) are the first principal component of nucleotide diversity and recombination rate of six *A. cerana* Lineage M groups (Japan, Qinghai, Bomi, Hainan, Taiwan and Central), respectively.

Genomic differentiation (*F*_ST_) values in low GC regions were always higher and recombination rate (rho) and nucleotide diversity (π) values lower than those in high GC regions (Supplementary Figure 5), thus supporting the linkage of low GC content with formation of genomic islands. We further characterized genomic islands (normalized *F*_ST_, Z-*F*_ST_ ≥ 2) over time and found a significantly lower GC content in genomic islands compared with genomic backgrounds (Z-*F*_ST_ < 2) (Supplementary Figure 6). Being consistent with the pattern found in low GC regions, recombination rate and nucleotide diversity in genomic islands were significantly lower than those in genomic backgrounds (Supplementary Figure 6). Comparisons of *d*_xy_ in genomic islands and backgrounds also varied over time, with lower-than-average *d*_xy_ values initially (t1 to t5) to higher-than-average at t6 and t7 (Supplementary Figure 6). This pattern supported the proposed model of genomic island formation changing from recurrent selection (i.e., high *F*_ST_ and low *d*_xy_) to divergent selection (i.e., high *F*_ST_ and high *d*_xy_) over deeper divergence.

### Contribution of recombination and AT-biased derived alleles to nucleotide diversity and group divergence

We further investigated the effects of genome architectures on nucleotide diversity. Rare, derived alleles (loci with derived allele frequency ≤ 1%, DAF_001) were mutations that arose recently and were used here as a proxy for mutation rate because natural selection had not yet worked on them (e.g., Coventry et al., 2010; Keinan and Clark, 2012). Correlation analysis showed that GC content was positively correlated with DAF_001 in both low and high GC regions across the genome (Figure 3). In addition, we found most of the mutations (∼80%) were derived A/T alleles from ancestral C/G ones (Supplementary Figure 7). This pattern supported the predominant role of GC content on newly generated mutations (i.e., AT-biased) across the genome (Hershberg and Petrov, 2010; Lynch 2007) with fewer mutations in low GC regions (Figure 3). This process could contribute to the low nucleotide diversity in low GC regions and therefore the presence of genomic islands (Supplementary Figure 2, 3, 5, 6).

To further distinguish the effects of different genomic architectures on genomic variation, we divided the genome into five regions (region 1 being low to region 5 being high GC) with equal-width intervals of GC content. We investigated the contribution of each genome architecture to nucleotide diversity using a regression analysis (Table 1). Four genomic architectures were included in the analysis: DAF_001, GC content, recombination PC1, and gene density. Principal component 1 (PC1) was extracted from recombination rate (explaining 74.91% of the total variance) and nucleotide diversity (explaining 86.02% of the total variance) across six groups in *A. cerana* Lineage M. Regression analysis across the five GC regions showed that the models explained between 40.3% (region 3) and 85.7% (region 1) of the variance in nucleotide diversity (Table 1). DAF_001 (i.e., proxy of mutation rate) had the strongest and significantly positive effect in all regions, with coefficients ranging from 0.4222 (region 4) to 0.6985 (region 3; all with p < 0.001). GC content showed a consistent negative association in each region, with the strongest value in region 4 (β = -0.5802, p < 0.001). The effect is pronounced because gBGC promotes the fixation of GC alleles, counteracting a genome-wide bias toward AT mutations (β=-2.341, p < 0.001). Recombination rate was generally positively associated, especially in region 1 (β = 0.4214, p < 0.001) and region 2 (β = 0.3638, p < 0.001), but negatively associated in region 3 (β = -0.1075, p < 0.01). Gene density was negatively associated in regions 1–5 (e.g., region 2: β = -0.2496, p < 0.001). Based on the relationships DAF_001 ∝ μ and ρ = 3Nl⍰c (where Nl⍰ is the effective population size, μ is the mutation rate, and c is the recombination rate, 3 is due to haplodiploid sex-determination system in honeybee), our models support important and independent roles for both mutation rate (proxied by DAF_001) and recombination rate (proxied by population scaled recombination rate) in shaping nucleotide diversity. Their contributions are particularly pronounced in low-GC regions (Table 1).

Following Wallberg et al. (2015), we further categorized mutation types in *A. cerana* groups using *A. mellifera* as the outgroup. There were four types of mutation S2S (strong to strong) (from C/G in *A. mellifera* to C/G in *A. cerana*), S2W (strong to weak) (from C/G to A/T), W2S (weak to strong) (from A/T to C/G), and W2W (weak to weak) (from A/T to A/T). In *A. cerana* lineage M (Figure 4A), relative site frequency spectrum of the four mutation types showed that an initially high S2W contributed by GC coupling with a fixation bias towards W2S mutations, consistent with a strong GC-biased gene conversion (gBGC) (Wallberg et al., 2015). We found, however, a higher percentage of nearly-fixed S2W mutations in the peripheral groups compared to the Central group and Lineage M, at both population genetic and phylogenetic levels (Figure 4B). This finding supports the contribution of fixed AT alleles to group divergence. But, the relative contribution of fixed AT alleles to group divergence varied among groups. For instance, the percentage differentiation of nearly fixed AT alleles is amplified by genetic drift (e.g., Japan vs. Lineage M Central Lineage) and divergence time (Lineage S vs. Lineage M, Figure 4B, C). In several groups (e.g., Qinghai, Hainan, Lineage S, Figure 4B, C), we found more fixation of AT-biased alleles in low GC regions (with lower AT mutations) compared with high GC regions (with higher AT mutations), suggesting the involvement of other mechanisms, such as local recombination and gBGC.

**Figure 4.**
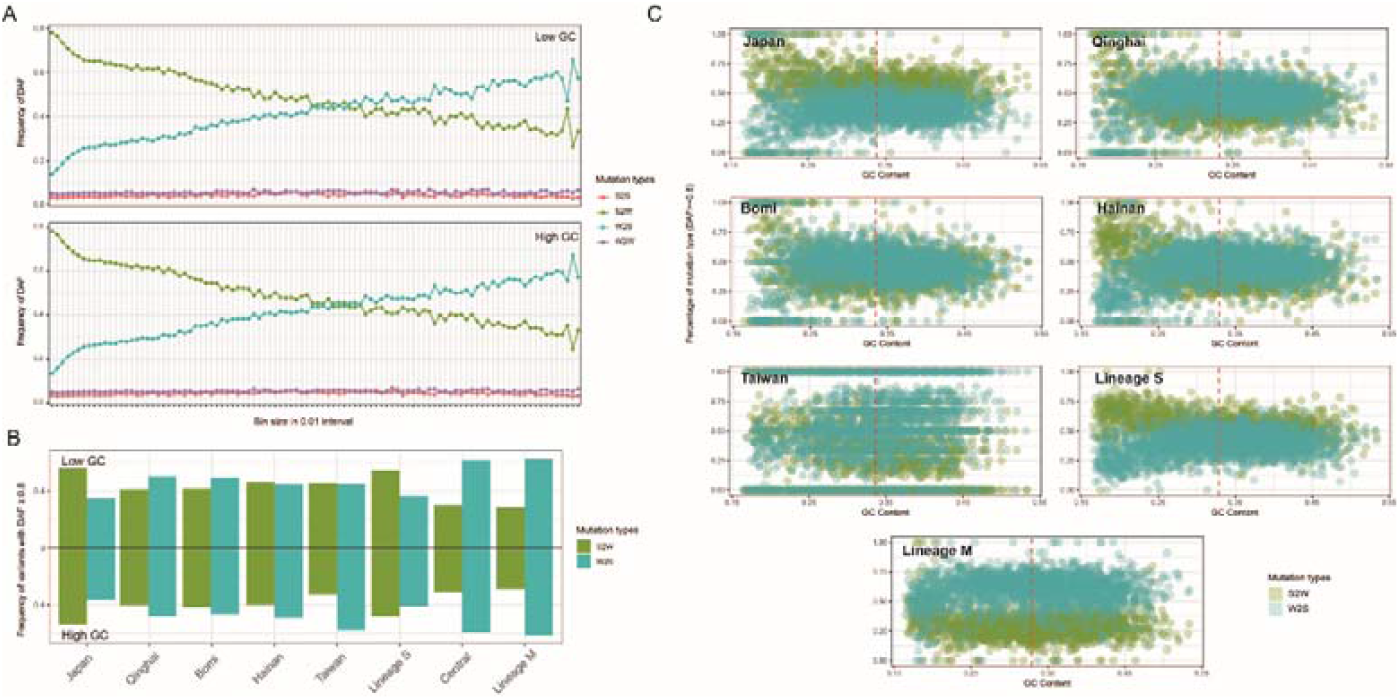
Variation of derived allele frequency (DAF) in low and high GC regions of different *A. cerana* groups. **(A)** The relative site frequency spectrum of four mutation types (bin size: 0.01, from 0-0.01 to 0.99-1) in *A. cerana* Lineage M inferred with *A. mellifera* as the outgroup. S2S: strong to strong (from C/G in *A. mellifera* to C/G in *A. cerana*), S2W: strong to weak (from C/G to A/T), W2S: weak to strong (from A/T to C/G), and W2W: weak to weak (from A/T to A/T). **(B)** Percentage differentiation of nearly fixed S2W and W2S alleles (DAF ≥ 0.8) in low and high GC regions of different *A. cerana* groups. Japan, Qinghai, Bomi, Hainan, Taiwan, Lineage S have higher percentage of S2W (C/G to A/T) type than Central and Lineage M, especially in low GC regions. **(C)** Percentage of nearly fixed S2W and W2S alleles in different *A. cerana* groups along the GC gradient. The red dash line represents the mean GC content.

### Profile of functional polymorphisms in low GC and high GC regions

Patterns of accumulated polymorphism (e.g., mutations) differed between low GC and high GC regions (Figure 5 A, B; Supplementary Figure 4E), with synonymous (dS) and non-synonymous (dN) substitution rates positively correlated with GC content in low GC regions and negatively correlated in high GC regions (Supplementary Figure 4). The differentiated pattern likely resulted from limited recombination and inefficient purifying selection in low GC regions (Figure 3; Supplementary Figure 5 and 8), which could have facilitated the accumulation of mutations, a pattern that could not be found in high GC regions with higher recombination (Figure 3). The consistently positive correlation between mutations and GC content in low GC regions indicated that these regions accumulated more mutations and divergence than high GC regions at deeper divergence (Figure 2A, B), although fewer mutations were found in these regions (Figure 3).

**Figure 5.**
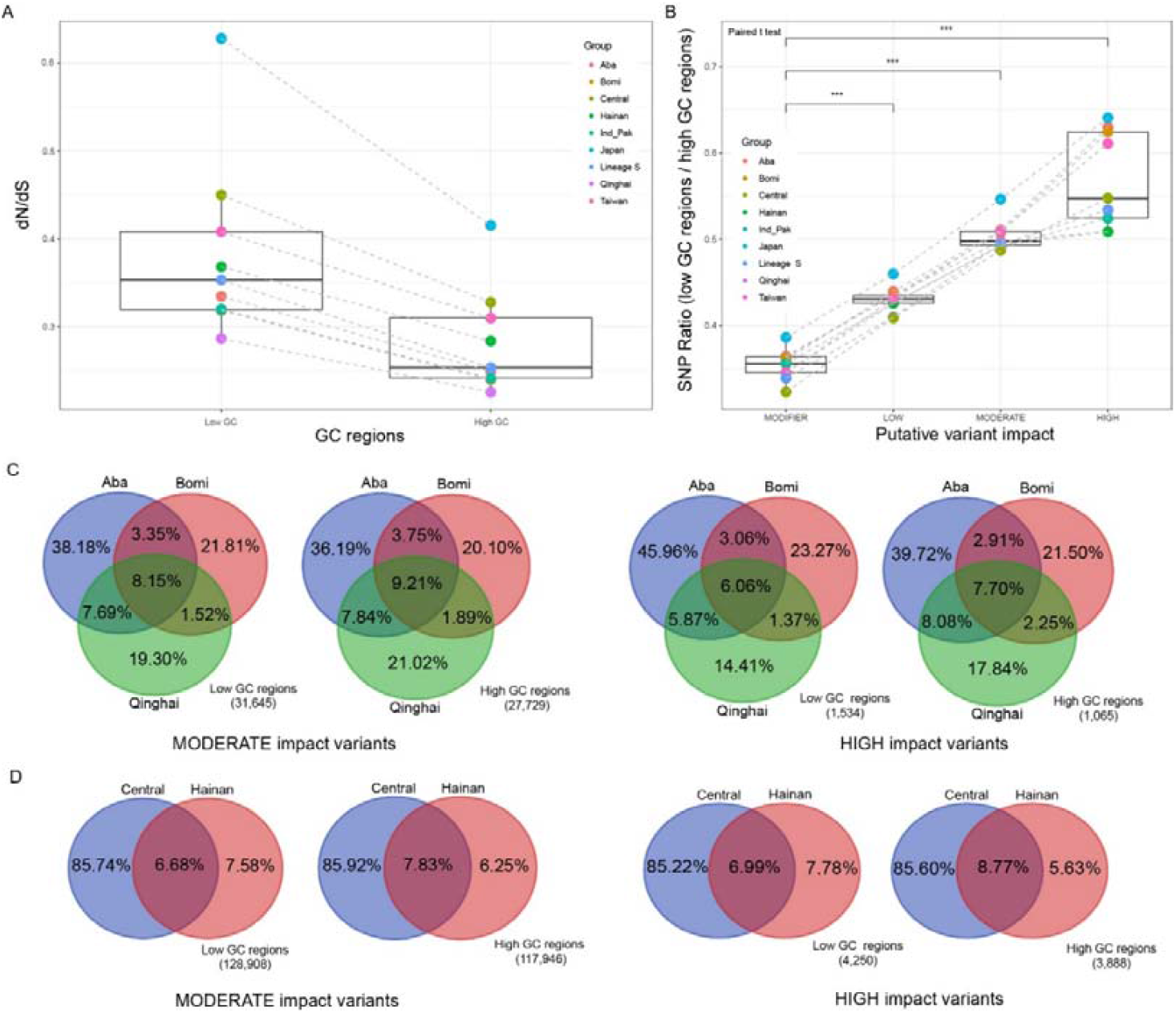
Profile of functional variants in high GC and low GC regions. **(A)** Higher dN/dS (ratio of nonsynonymous over synonymous mutations) in low GC regions than high GC regions (Paired t test, p < 0.001). **(B)** Higher ratio of putatively deleterious mutations than neutral mutants. Paired t tests were conducted between ratio of different mutant types. *** indicates p < 0.001. Types of putative variant impact could refer to SNPeff (Cingolani et al., 2012); **(C)** Consistent pattern of more group-specific mutation load than the shared in both high GC and low GC regions of isolated groups. Aba, Bomi and Qinghai represent three independent colonization to Qinghai-Tibet Plateau and are with no admixture with each other (Figure 1). **(D)** Ratio of group-specific and shared mutation load differentiated in low GC and high GC regions in group pair with gene flow. Group pair Central vs. Hainan, as an example.

We calculated dN/dS and found significantly higher values in low GC regions than in high GC regions (Figure 5A). We further divided the mutation load into variants with different impacts (MODIFIER, LOW, MODERATE, and HIGH) and calculated the ratio of SNP numbers in low GC over high GC regions. We found significantly higher ratios of non-neutral variants (LOW, MODERATE, and HIGH) than neutral ones (MODIFIER) in all defined groups (Figure 5B), which indicated low GC regions could accumulate more impact SNPs than high GC regions due to inefficient purifying selection. This pattern supported the idea that the genome architectures (i.e. restricted recombination) was involved in mutation accumulation and divergence in low GC regions of *A. cerana* groups due to Hill-Robertson interference (Hill and Robertson, 1966).

We further investigated the ratio of polymorphisms that was group-specific or shared among different groups in low GC and high GC regions. With three groups (Bomi, Aba and Qinghai) that independently colonized highland habitats and had no genetic admixture (Figure 1), we determined that most of the polymorphisms (e.g. mutation load) were unique to each group and showed no percentage difference in high GC and low GC regions (Figure 5C). For example, in MODERATE impact polymorphism, Aba had 38.18% (12,082) and 19.19% (6,073) of unique and shared variants in Low GC regions, and 36.19% (10,035) and 20.8% (5,768) in High GC regions. We further investigated group-specific and shared polymorphisms in lineage pairs with genetic admixture. With Central and Hainan group pair as an example, we found more group-specific than shared polymorphism only in low GC regions (Figure 5D). For MODERATE impact polymorphism, Hainan had 7.58% (9,771) and 6.68% (8,611) unique and shared variants in low GC regions, corresponding to 6.25% (7,372) and 7.83% (9,235) in high GC regions.

### Reconstruction of A. cerana nuclear phylogeny

After characterizing the landscape of genomic variation in high GC and low GC regions, we reconstructed the phylogeny of *A. cerana* based on nuclear polymorphism in low GC regions with more group-specific polymorphisms (Figure 5D). We reconstructed a nuclear topology similar to the mitochondrial phylogeny (Figure 6 and Supplementary Figure 9, mitochondrial phylogeny from Qiu et al. 2023), with individuals from Taiwan and Hainan being basal groups of *A. cerana* Lineage M. The phylogeny based on polymorphism in high GC regions was influenced by gene flow, with admixed groups clustering together (Supplementary Figure 10). Individuals showing hybrid ancestry were always situated between groups (e.g. Central vs. derived group, Figure 1 and Supplementary Figure 10) and had higher polymorphism than the individuals within the derived groups (Supplementary Figure 10).

**Figure 6.**
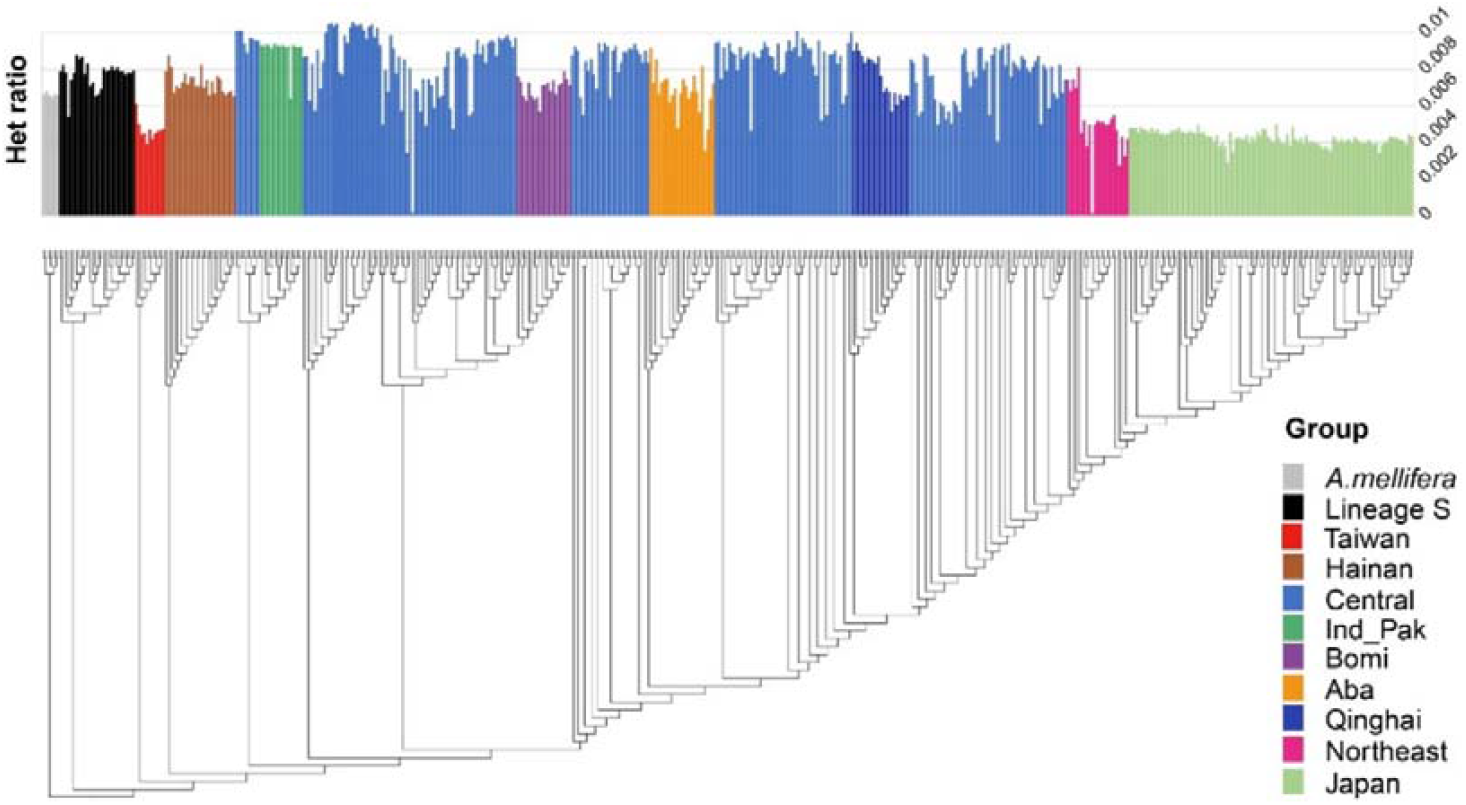
Reconstruction of *Apis cerana* phylogeny based on nuclear polymorphisms in low GC regions. Bootstrap values and individual IDs could be referred to phylogeny in Supplementary Figure 8; Het ratio indicates number of heterozygous sites/number of homozygous sites in each individual.

## Discussion

The formation of heterozygous differentiation landscape is suggested to result from the combined effects of external evolutionary processes and genome architectures (Ravinet et al. 2017). Here, we used 499 eastern honeybee genomes along the speciation continuum to investigate the influences of GC content on mutation accumulation and group differentiation by conducting correlation analysis of population genetic parameters and genomic architectures, characterization of different genomic regions, and regression analysis of genome architectures on genomic variation. We found that the local GC content could elevate regional mutations because GC to AT mutations accounted for around 80% of the newly generated alleles (Supplementary Figure 7). This however could be counteracted by GC-biased gene conversion (gBGC) that lead to fixation of GC alleles.. We also found local GC content was positively correlated with recombination rate that further interplayed with evolutionary forces and shaped mutation accumulation and group differentiation across the genome.

Over the diverging groups, genomic islands have significantly lower GC content and recombination compared with background regions (Supplementary Figure 4). This is consistent with fewer mutations (i.e., lower GC) and limited recombination that leads to lower nucleotide diversity and elevated differentiation (Cruickshank and Hahn 2014). In addition, genomic islands with reduced nucleotide diversity were shared across diverging groups of *A. cerana* (Supplementary Figure 2, 3), suggesting the important role of common genome architectures (Barroso and Dutheil 2023).

The lower-than-average divergence (*d*_xy_) in GC regions at population genetic timescales (within species) to higher-than-average *d*_xy_ at phylogenetic levels (between species) corresponds to the suggested evolution models transformed from “sweep-before-differentiation” and background selection (i.e. recurrent selection, Charlesworth et al. 1993; Irwin et al. 2016; Ravinet et al. 2017), to divergent selection with gene flow (Feder et al. 2012; Nosil et al. 2009; Wolf and Ellegren 2017) or sorting of ancient polymorphisms (Guerrero and Hahn 2017; Han et al. 2017; Noor and Bennett 2009; Wolf and Ellegren 2017). Correlations between *d*_xy_ and recombination rate (rho) in low GC regions were negative in population genetic timescale but positive in phylogenetic timescale. A similar trend was also found in the correlations between *d*_xy_ and *F*_ST_. These results align with previously proposed models of genetic island formation, which suggest that initial formation involves recurrent selection in regions with low recombination (Ravinet et al. 2017). In contrast, divergent selection due to the breakdown of linkage occurs over deeper divergence (Cutter and Choi 2010; Noor and Bennett 2009). Few studies have reported a similar pattern (increased *d*_xy_ values in genomic islands along the speciation continuum, e.g., Liu et al. 2022, and shift pattern of parameter correlations) likely because few have analyzed group pairs at both population genetic (within species) and phylogenetic (between species) timescales in a single system. Many studies report on one dominant model in genomic island formation, with the contribution of external processes and genomic architectures. For example, recurrent selection is reported to be involved in the formation of genomic islands in population pairs of Green-backed Tit (*Parus monticolus*) (Jiang et al. 2023). Zhou et al. (2024) indicate that divergent selection contributes to regionally elevated divergence and genomic islands in incipient species of plover (*Charadrius alexandrines and C. dealbatus*). However, reductions in recombination were needed in the formation of these genomic islands in both cases (Jiang et al. 2023; Zhou et al. 2024).

Synonymous (dS) and non-synonymous (dN) polymorphisms, and mutation load (dN/dS) in both low GC and high GC regions were also analyzed. The results showed higher mutation loads (i.e. higher dN and dN/dS ratio) in low GC regions than high GC regions, suggesting inefficient natural selection (Arbuthnott and Rundle 2012; Bertorelle et al. 2022; Leroy et al. 2021) in these regions compared with the rest of the genome. It is worth noting that this pattern was not only in peripheral groups with lower genome-wide nucleotide diversity (e.g., Japan group) where it might be attributed to genetic drift (Bertorelle et al. 2022). Instead, it was observed across all *A. cerana* groups including *A. cerana* Lineage M Central group and Lineage S. This broader finding supports the continuous effects of restricted recombination leading to the accumulation of neutral polymorphisms and mutation load due to Hill-Robertson interference (Hill and Robertson, 1966) in low GC regions. The relative contribution of external evolutionary processes and genome architectures to regional variation and differentiation thus needs to be reconsidered. The reduced nucleotide diversity of the genomic islands may not result from natural selection (Feder and Nosil 2009; Turner and Hahn 2010), genetic drift, or both (Burri 2021). It could be independently contributed by local genomic architectures, where lower mutation rate governs the accumulations of less random mutations (Castellano et al. 2020), and reduced recombination protects low GC regions from natural selection (Hill and Robertson, 1966; Leroy et al. 2021) and gene flow (Wong and Filatov 2023) along the speciation continuum. Therefore, the genomic islands identified in *A. cerana* groups—whether at the population genetic timescale (characterized by high *F*_ST_ and low *d*_xy_) or the phylogenetic timescale (characterized by high *F*_ST_ and high *d*_xy_)—could be largely explained by the influence of genome architectures (such as low mutation and recombination rates) rather than by specific evolutionary processes like recurrent or divergent selection.

In contrast to low GC regions, high recombination in high GC regions facilitate natural selection (Roze 2021) and genetic admixture (Wong and Filatov 2023). This leads to lower mutation load and reduces differentiation in these regions at group pairs of both intraspecific and interspecific levels. Therefore, admixed ancestry in high GC regions could blur nuclear phylogeny of two diverging groups in gene-flow scenario and cause conflict between nuclear and cytoplasmic DNA topologies of *A. cerana* (e.g., Ji et al. 2020; Qiu et al. 2023). The conflict, however, could be reconciled by polymorphisms in low GC regions with restricted recombination (Pease and Hahn 2013). These findings supported differentiated recombination in high and low GC regions related to regional nucleotide diversity and accumulation of mutation loads (e.g., lead to lineage divergence) (Barroso and Dutheil 2023).

Our results suggested that genome architectures such as regional mutation and recombination could shape mutation accumulation (e.g., lower nucleotide diversity in low GC regions) and group divergence (e.g., more group-specific alleles in low GC regions) at different genome regions along divergent groups. Genomic studies often reveal correlations between variation and divergence in evolving groups across various species (e.g., Cuevas et al., 2022; Vijay et al., 2016). These patterns are likely influenced by conserved genomic architectures, such as recombination (Burri et al., 2015; Renaut et al., 2014; Shang et al., 2023). The influence of local mutations and gBGC in nucleotide diversity and genetic differentiation has received less investigation (Boman et al., 2021). Due to the significant effects of regional A/T biased mutation, gBGC, and recombination on nucleotide diversity and fixation of alleles, we would suggest considering the effects of local genome architectures in future analyses related to the formation of heterozygous differentiation landscapes along diverging groups and identification of adaptive loci based on summary parameters (e.g., *F*_ST_ and *d*_xy_).

Genome architectures, such as variation of recombination rate (Booker et al., 2020), have been widely reported to produce pseudo positives that could not be easily distinguished from those generated in non-neutral processes in genome-scan methods (e.g., Southcott & Kronforst, 2017). When explaining signals from genome-scan methods, caution is advised, and stricter methods should be applied when isolating adaptation and speciation loci. Several parameters have been introduced to refine the analysis of population differentiation statistics and minimize false positives. For *F*_ST_ statistic, Δ*F*_ST_ (Burri, 2017) and Population Branch Statistic (PBS) (Yi et al., 2010), for example, have been proposed. These approaches leverage one or several reference populations to reduce pseudo positive signals. When calculating dxy or other linkage-based parameters like the haplotype homozygosity statistic (Garud et al., 2015), the use of a control population can help eliminate false positives, thus reducing burden in further functional validation efforts.

## Materials & Methods

### Sequencing data processing, SNP calling, and variant filtering

A total of 507 *Apis cerana* workers were initially sampled across its distribution range from previous studies, including individuals from China (331), India (14), Japan (105), Malaysia (26), Myanmar (2), North Korea (1), Pakistan (5), Singapore (1), South Korea (6), Thailand (11) and Viet Nam (5) (Chen et al. 2018; Ji et al. 2020; Qiu et al. 2023; Wakamiya et al. 2023). Six *A. mellifera* individuals (Harpur et al. 2014) were used as the outgroup individuals. After preprocessing, eight *A. cerana* individuals with elevated percentage of missing sites (≥20%) or with unknown sampling sites were removed, resulting in a total of 505 individuals (499 *A. cerana* and six *A. mellifera* individuals) to be used for the analysis (Supplementary Table 1).

All sequencing reads were downloaded from National Center for Biotechnology Information (NCBI) Sequence Read Archive (SRA) (PRJNA592293, PRJNA418874, PRJNA870246, PRJNA216922), or DNA Data Bank of Japan (DDBJ) database (PRJDB14080). Clean reads were trimmed and filtered using Trimmomatic (v 0.39, Bolger et al. 2014). Reads with a minimum length of 50bp were kept for analysis. We further individually mapped clean reads to the *A. cerana* genome (GCF_001442555.1, Park et al. 2015) by using bwa (v 0.7.17, Li 2013). Duplicates were identified by Picard (v 2.24.0, http://broadinstitute.github.io/picard/). A standard pipeline of variant calling based on Genome Analysis Toolkit (GATK 4.2.4.0, McKenna et al. 2010) was applied with HaplotypeCaller for indel realignment and individual variant calling, and GenotypeGVCFs for joint variant genotyping in all samples with “--include-non-variant-sites and --heterozygosity 0.001”. We further performed hard filtering based on suggested parameters in GATK by using VariantFiltration command (--filter-expression “SOR > 3.000”, --filter-expression “FS > 60.000”, --filter-expression “MQRankSum < -12.500”, --filter-expression “ReadPosRankSum < -8.000”, --filter-expression “ReadPosRankSum > 8.000”, --filter-expression “MQ < 40.00”, --filter-expression “QD < 2.00”). In addition, hard filtering was conducted with Vcftools (v 0.1.16, Danecek et al. 2011) using “—minDP 3 --min-meanDP 3 --max-meanDP 60 --minQ 30 --max-alleles 2 --max-missing 0.5 --remove-indels --remove-filtered-all” to generate a high-quality dataset. This dataset, including non-variant sites, was used for calculation of population genetic parameters, including nucleotide diversity (π) and divergence (*d*_xy_).

We also used Vcftools (v 0.1.16, Danecek et al. 2011) in SNP filtering when investigating genetic structure and reconstructing phylogeny. In genetic structure analysis, bi-allelic SNPs with a maximum missing value of 20% and a minor allele frequency of 0.05 were extracted, after which one SNP from a window of 20-kb DNA sequence was kept for analysis. We used a 20-kb window for thinning to remove linkage of polymorphic sites in genetic structure inference. For window phylogeny reconstruction with maximum likelihood method (see **2.5 Nuclear phylogeny reconstruction**), we kept all bi-allelic SNPs in the entire dataset for the analysis.

### Genetic structure and population genetic parameter calculation

We used ADMIXTURE (v 1.3.0, Alexander et al. 2009) for genetic structure analyses and performed each K from 2 to 20 with 30 replicates. The best K value was identified according to cross-entropy criterion. The outgroup individuals (A. mellifera) were excluded for the genetic clustering analysis. To characterize the environmental conditions for each sample, we obtained monthly climatic variables (e.g., temperatures, precipitation) and water balance for global terrestrial surfaces from 1996 to 2015 (high-spatial resolution, 1/24°, ∼4-km) (Abatzoglou et al. 2018). We then performed a Principal Component Analysis (PCA) with twelve monthly averages of the twelve variables (a total of 144 monthly variables, Supplementary Table 1) to investigate environmental heterogeneity of the sampling locations.

To have an unbiased calculation of π and *d*_xy_, we used the dataset with variant and non-variant sites. We calculated π, *d*_xy_, *F*_ST_ in a 100 kb window size within and among our identified groups using Python scripts downloaded from github.com/simonhmartin/genomics_general. We removed all individuals that had more than 20% admixed ancestry from the other group, resulting in 24 (Aba), 20 (Bomi), 224 Central, 26 (Hainan), 16 (Ind_Pak), 105 (Japan), 27 (Lineage S), 23 (Northeast), 21 (Qinghai) and 11 (Taiwan). Six *A. mellifera* (i.e., Mel, outgroup) were also included in parameter calculation. Windows with less than 1000 genotyped sites (variant and invariant) were removed from the analysis. Regional GC-content and coding sequence percentage were also calculated in 100 kb window based on the *A. cerana* genome sequence (GCF_001442555.1). For each group, calculation of recombination rate was conducted by using FastEPRR (v 2.0, Gao et al. 2016). Dataset with variant sites was extracted with Vcftools (v 0.1.16, Danecek et al. 2011) and phased using beagle (v 5.4, Browning et al., 2021) with default settings. We then calculated population-scaled recombination rate of each group in FastEPRR in a sliding window size of 100kb.

Within *A. cerana* Lineage M, we further estimated divergence times of five groups (Bomi, Hainan, Japan, Qinghai and Taiwan) with the Central group based on coalescent theory in two-two-outgroup (TTo) method (Sjödin et al., 2021), which is relatively robust to migration and could alleviate biases due to drastic ancestral population size change (Sjödin et al., 2021). The ancestral state of the sites was inferred with A. mellifera, and *A. cerana* Lineage S was used as the outgroup in TTo method. Since Central group of *A. cerana* Lineage M showed divergence in Southern part and Northern part of the distributed region, we calculated and checked the divergence time of each group with Central group using individuals from Southern and Northern region of Central group. The mutation rate μ equals to 5.27 x 10^-9^ per base pair following Wallberg et al. (2014) and generation time g to be 1 year (Ji et al., 2020).

### Characterization of summary statistics in different regions

The pilot analysis and visualization of genome-wide parameters showed distinct pattern of differentiation in high GC and low GC regions (Figure 2A, Supplementary Figure 1). Therefore, we divided the genome-wide windows into high GC (higher than average GC content) and low GC (lower or equal to average GC content) categories. Difference in parameter values in high GC and low GC windows were compared and tested based on Wilcoxon rank sum test in ggsignif R package (v 0.6.4, Ahlmann-Eltze and Patil 2021). We further standardized per-window *F*_ST_ in each group pair to a Z score (Han et al. 2017) based on the formula: Z-*F*_ST_ = (window *F*_ST_ − mean *F*_ST_) / standard deviation *F*_ST_. We then compared the parameter values in the genomic islands (windows with Z-*F*_ST_ ≥2 from each group pair) and background regions (windows with Z-*F*_ST_ < 2) using a Wilcoxon rank sum test.

We conducted correlations of summary statistics to investigate the effects of different evolutionary forces on formation of heterozygous genomic landscape. The Pearson’s correlation coefficient for intragroup summary statistics between groups (e.g. π_i_ vs. π_j_), within groups (e.g. π_i_ vs. rho_i_), and intergroup statistics of group pairs (e.g. π_i_ vs. F_STij_) were calculated and performed in R package corrplot (v 0.94, Wei and Simko 2021). The correlation analysis for each group pair was also performed in high GC and low GC regions, respectively.

To investigate the effects of genome architectures on polymorphisms, we grouped all individuals from *A. cerana* Lineage M and performed correlations between GC content and recombination rate, gene density, and polymorphism of *A. cerana* Lineage M, including rare derived alleles (with derived allele frequency ≤ 0.01, DAF_001), and nucleotide diversity (π). We extracted principal component 1 (PC1) for recombination rate (explained 74.91% variance) and nucleotide diversity (explained 86.02% variance) from six *A. cerana* Lineage M groups for analysis. In order to capture variation across the genome instead of groups, we normalized each parameter (i.e., recombination rate or nucleotide diversity) in each group before analysis for PC1 extraction. To identify variables contributing to nucleotide diversity (nucleotide diversity PC1), we further conducted regression analysis with genome architectures (DAF_001, GC content, recombination PC1, and gene density) in high-GC and low-GC regions, as well as in five divided regions based on equal-width intervals of GC content.

The allele state (ancestral or derived) of each locus in *A. cerana* was inferred with *A. mellifera* as the outgroup. The derived allele frequency of each locus in each *A. cerana* group was then calculated with Vcftools (v 0.1.16, Danecek et al. 2011). With *A. mellifera* as the outgroup, we further sorted and counted the number of four mutation types in each *A. cerana* groups following Wallberg et al. (2014), including S2S: strong to strong (from C/G in *A. mellifera* to C/G in *A. cerana*), S2W: strong to weak (from C/G to A/T), W2S: weak to strong (from A/T to C/G), and W2W: weak to weak (from A/T to A/T). We then investigated the relative site frequency spectrum of four mutation types in each *A. cerana* group and along the GC gradient.

### Investigation of mutation load

Synonymous and non-synonymous mutations in high GC and low GC regions in each group were investigated. Bi-allelic SNPs were extracted from each group and functionally annotated with SNPeff (v 5.1d, Cingolani et al. 2012), a variant annotation and effect prediction tool. In each group, we then calculated dN/dS (non-synonymous / synonymous SNPs) ratio that represents magnitude of mutation load. In addition, numbers of SNPs in low GC regions divided by high GC regions were calculated in each functional group (MODIFIER, LOW, MODERATE, and HIGH) in the defined groups. To further characterize whether the polymorphisms were shared or group-specific, mutation load in three highland groups (i.e., Aba, Bomi, and Qinghai) that colonized Qinghai-Tibet Plateau independently and with no admixture with each other, was investigated. As a comparison, mutation loads in group pair with admixture (i.e., Central and Hainan) were further investigated to estimate the effects of gene flow on group-specific and shared variants in low GC as well as high GC regions.

### Nuclear phylogeny reconstruction

For phylogeny reconstruction, we divided the genome into multiple 200-kb windows, and reconstructed window phylogeny with maximum likelihood method by using IQ-TREE 2 (v 2.2.2.7, Minh et al. 2020). In model testing, we employed +ASC model to include ascertainment bias correction as SNP data did not contain constant sites (Minh et al. 2020). We also appended -bnni to the regular UFBoot command to reduce the risk of overestimating branch support according to Hoang et al. (2018). Since genomic regions with high recombination rate and low recombination rate showed differentiated efficiency in tree reconstruction (e.g., Pease and Hahn 2013), and our analysis showed distinct pattern of recombination and mutation rates in high GC and low GC regions (Figure 3, Supplementary Figure 4, 5), we divided genomic windows into high GC (above genome-wide average GC content) and low GC (below genome-wide average GC content) categories, and generated the species tree in ASTRAL III (v 5.7.1, Zhang et al. 2018) based on window trees in each category.

## Supporting information

Shell scripts for analusis

## Acknowledgments

We would like to thank the Information Technology Services in HKU and in NUS for offering research computing facilities in the computations of this project. We acknowledge the assistance of AI-Know (https://ai-know.nus.edu.sg/ai-chat) and DeepSeek (https://chat.deepseek.com/) in writing scripts used during the analysis of this research. We also thank Dr. Per Sjödin’s help in divergence time calculation with TTo method.

## Data and code availability

All sequencing data could be retrieved from public data base. Software and packages for analysis in this project are all available online. Shell scripts were attached in supplementary files.

## Author contributions

Conceptualization, F.S.K.; methodology and analysis, F.S.K.; Writing, F.S.K. and L.V.

---

**Supplementary Table 1.** Sample and sequencing information of *Apis cerana*.

**Supplementary Table 2.** Divergence time estimation between *A. cerana* Central group with other seven groups.

**Supplementary Figure 1**. Landscape of genomic divergence (dxy) over time varied along with regional GC content. Central vs. Japan (t1), Central vs. Qinghai (t2), Central vs. Bomi (t3), Central vs. Hainan (t4), Central vs. Taiwan (t5), Central vs. Lineage S (t6), and Central vs. Mel (t7).

**Supplementary Figure 2**. Landscape of genomic variation (π) varied along with regional GC content.

**Supplementary Figure 3**. Landscape of genomic differentiation (*F*_ST_) over time varied along with regional GC content. Central vs. Japan (t1), Central vs. Bomi (t3), and Central vs. Lineage S (t6).

**Supplementary Figure 4**. Correlation coefficients of genome-wide parameters over divergence time.

**Supplementary Figure 5**. Characterization of low GC and high GC regions over divergence time. **(A)** *F*_ST_; **(B)** recombination (rho); **(C)** nucleotide diversity (π).

**Supplementary Figure 6**. Characterization of genomic islands and background regions over divergence time. **(A)** *d*_xy_; **(B)** recombination (rho); **(C)** nucleotide diversity (π); **(D)** dN/dS; **(E)** GC content.

**Supplementary Figure 7**. Summary of mutation types in all derived alleles. Each panel represents loci within a specific range of derived allele frequency. The plot shows a high percentage of derived A/T alleles from ancestral C/G alleles (i.e. C/G_2_A/T), especially in loci with rare, derived alleles.

**Supplementary Figure 8**. Site frequency spectrum in different regions. Rare/common ratio of SNPs in the inset plot suggest more efficient selection in high GC regions than in low GC regions.

**Supplementary Figure 9**. Reconstruction of *Apis cerana* phylogeny based on nuclear polymorphisms in low GC regions. Bootstrap values over 0.7 were presented.

**Supplementary Figure 10**. Reconstruction of *Apis cerana* phylogeny based on nuclear polymorphisms in high GC regions. Het/Hom indicate ratio of homozygous *vs*. heterozygous sites in an individual. The hybrid individuals referred to genetic structure in Figure 1.

