## Supplementary Figures for "Contribution of local recombination and AT-biased mutations to differentiated region formation in *Apis cerana*"

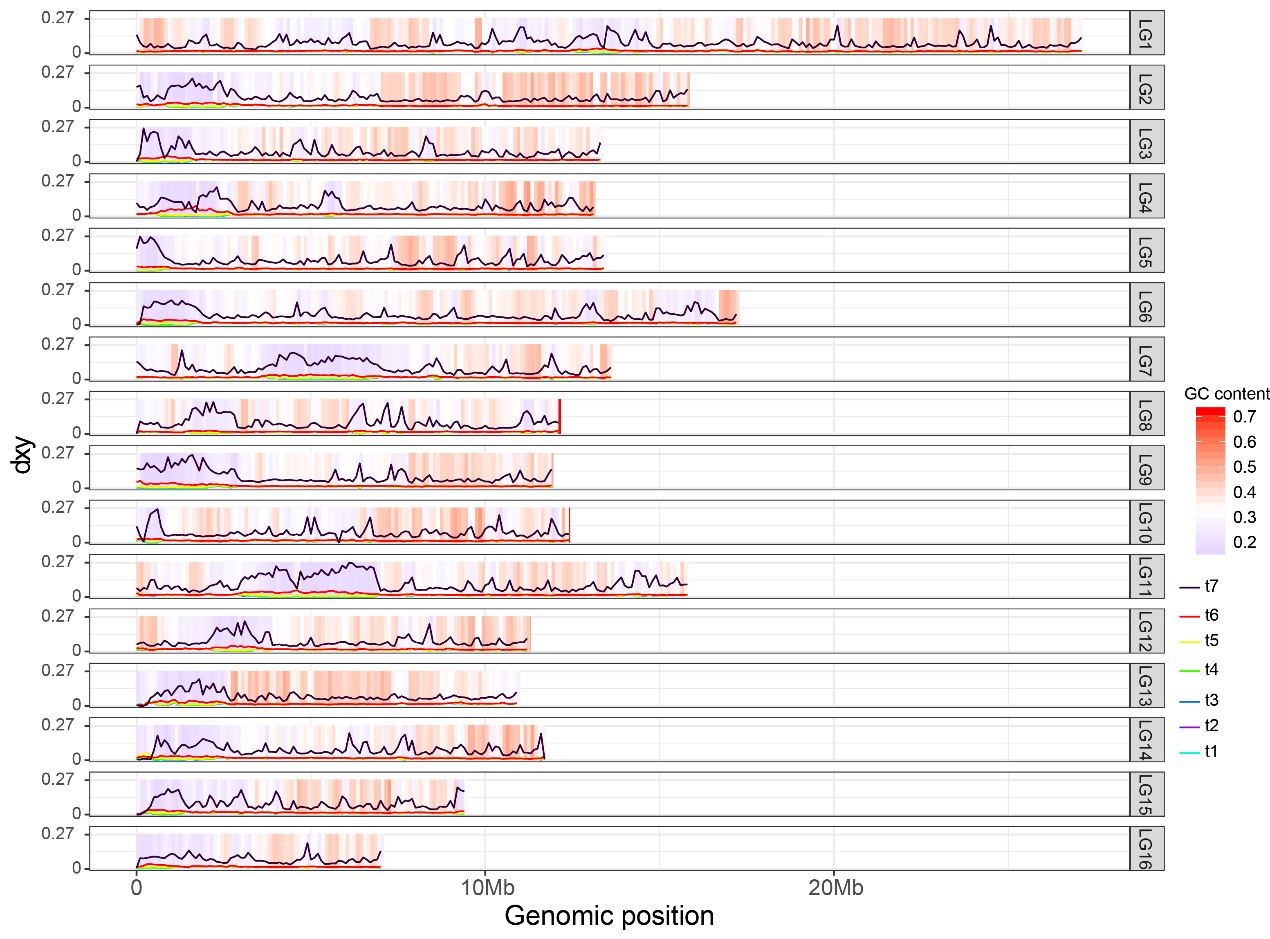

**Supplementary Figure 1.** Genomic landscape of lineage divergence (*d*_xy_) over time varied along with regional GC content. Central vs. Japan (t1), Central vs. Qinghai (t2), Central vs. Bomi (t3), Central vs. Hainan (t4), Central vs. Taiwan (t5), Central vs. Lineage S (t6), and Central vs. Mel (t7).

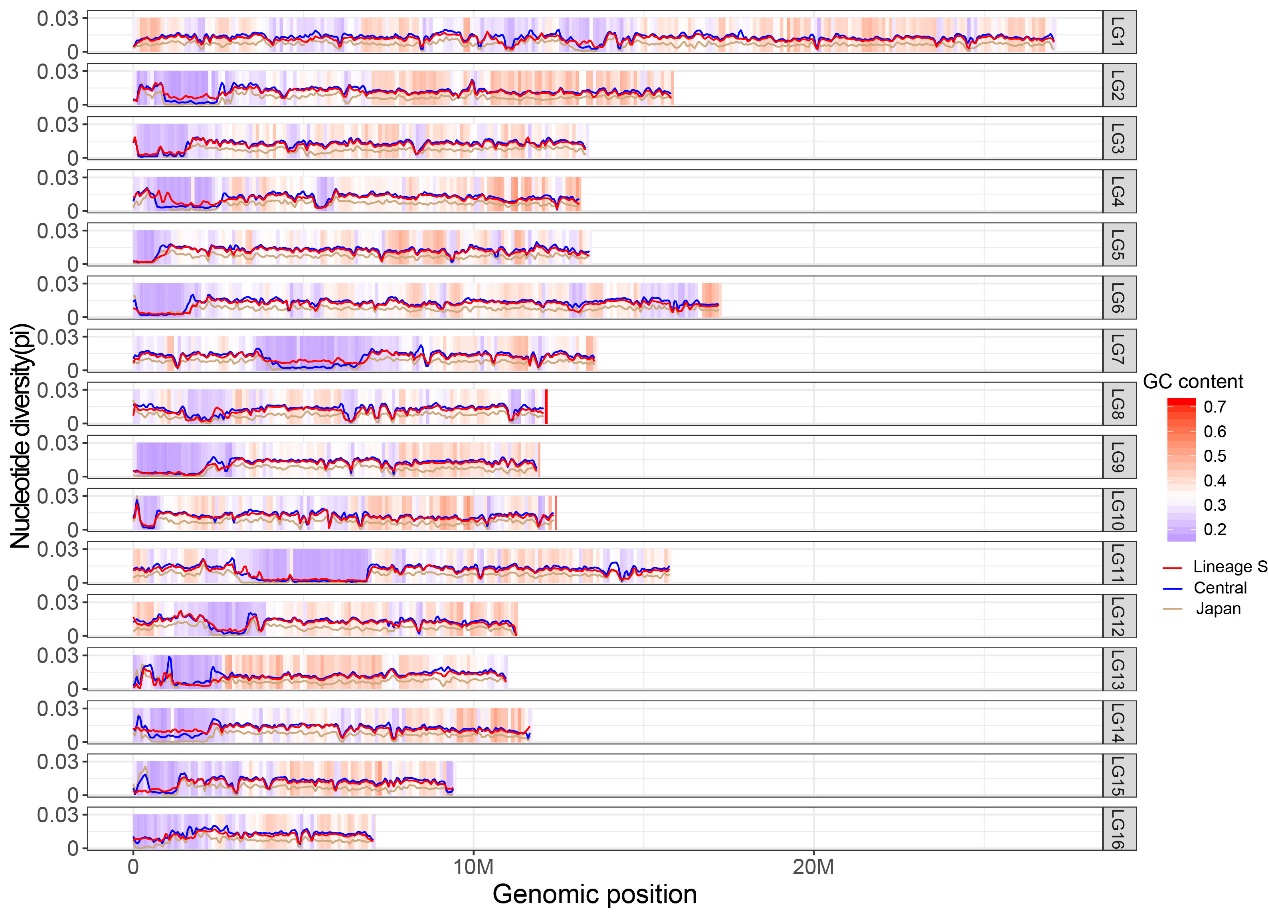

**Supplementary Figure 2.** Landscape of genomic variation varied along with regional GC content.

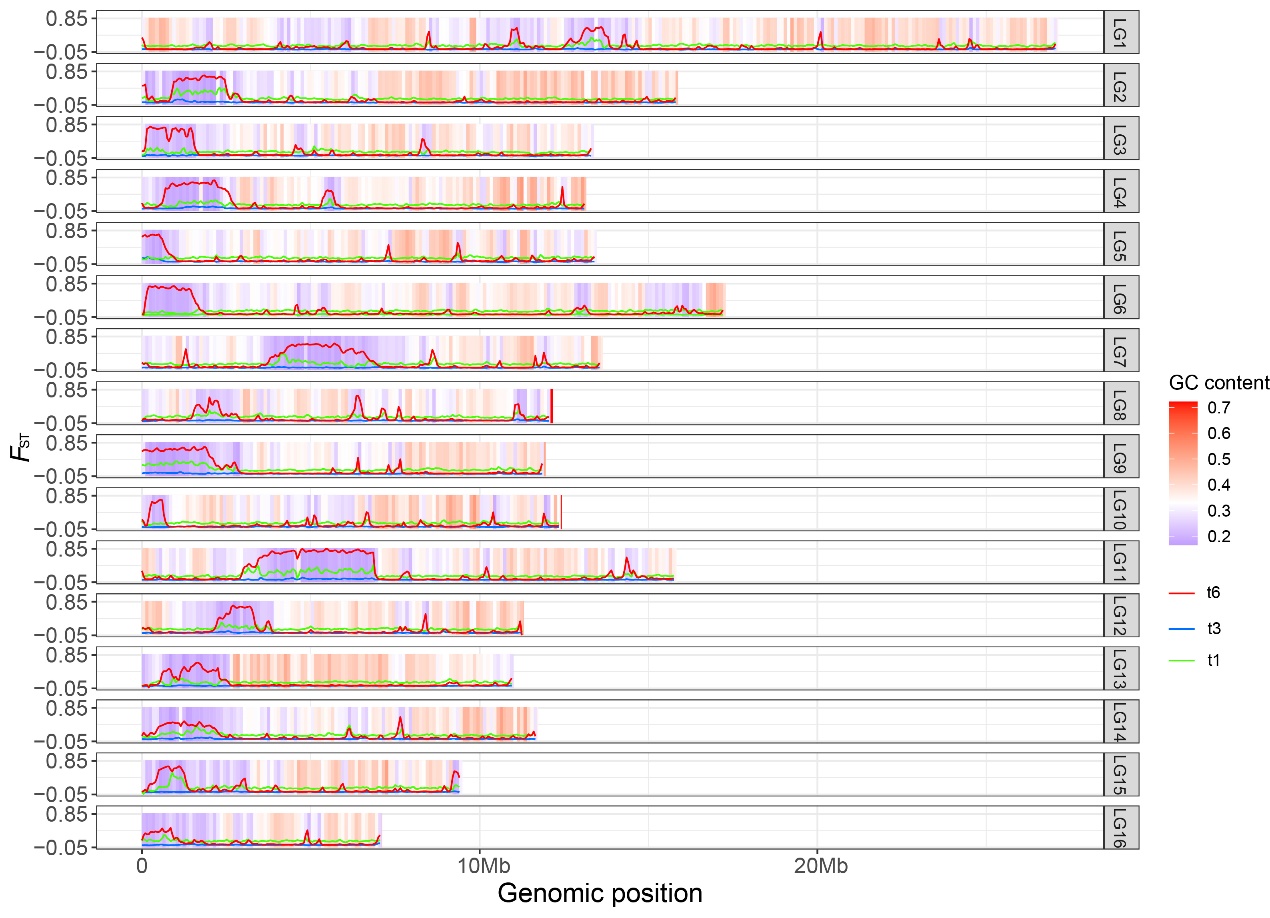

**Supplementary Figure 3.** Landscape of genomic differentiation (*F*_ST_) over time varied along with regional GC content. Central vs*.* Japan (t1), Central vs*.* Bomi (t3), and Central vs*.* Lineage S (t6).

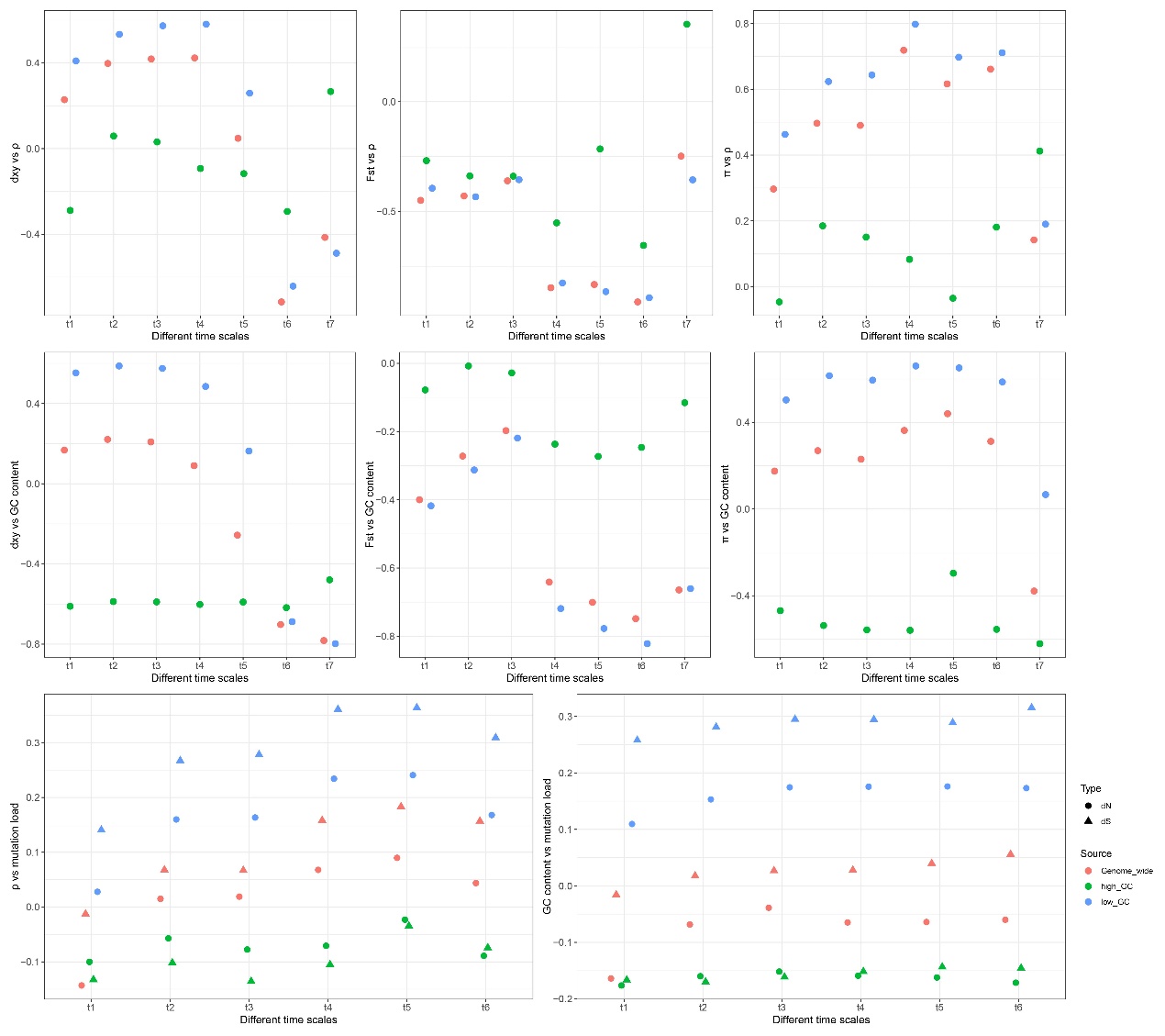

**Supplementary Figure 4.** Correlation coefficients of genome-wide parameters over divergence time.

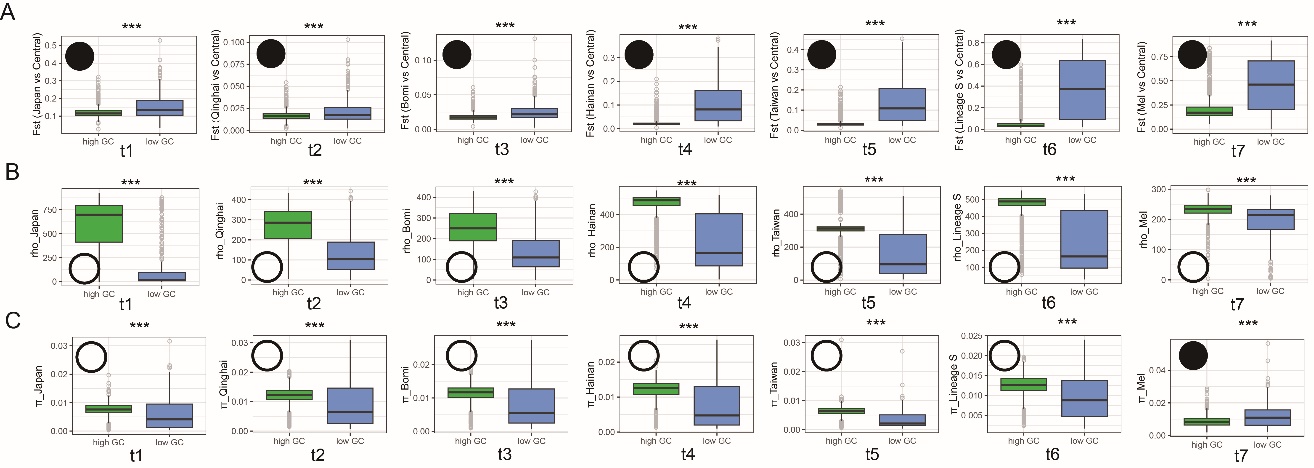

**Supplementary Figure 5.** Characterization of low GC and high GC regions over divergence time. **(A)** *F*_ST_; **(B)** recombination (rho); **(C)** nucleotide diversity (π). Wilcoxon rank sum test was conducted with values between low GC and high GC regions. Empty circles represent significantly lower values in low GC than high GC regions, while black circles indicate low GC regions with higher values.

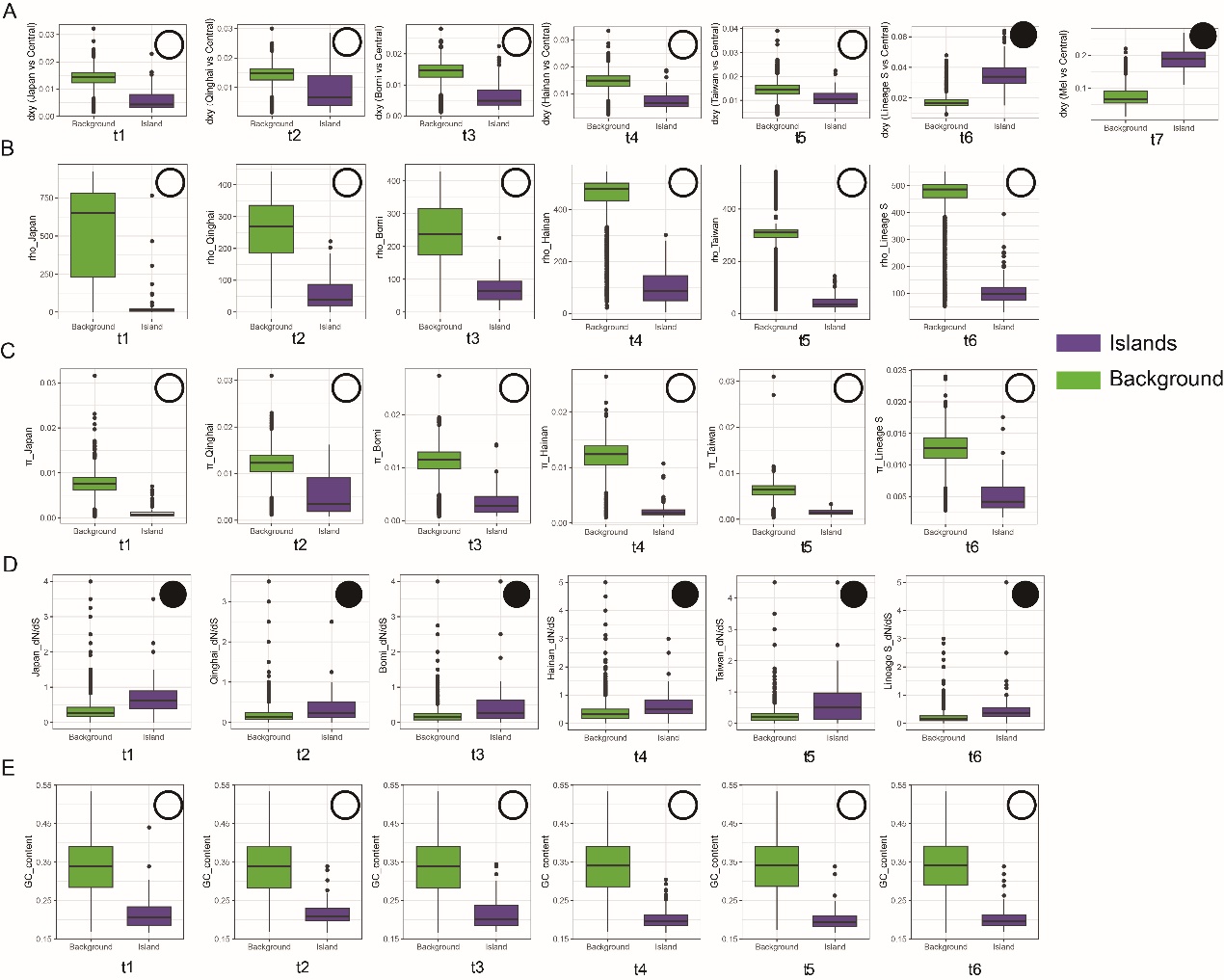

**Supplementary Figure 6.** Characterization of genomic islands and background regions over divergence time. **(A)** *d*_xy_; **(B)** recombination (rho); **(C)** nucleotide diversity (π); **(D)** dN/dS; **(E)** GC content. Wilcoxon rank sum test was conducted with values between genomic islands and background regions. Empty circles represent significantly lower values in genomic islands than background regions, while black circles indicate genomic islands with higher values.

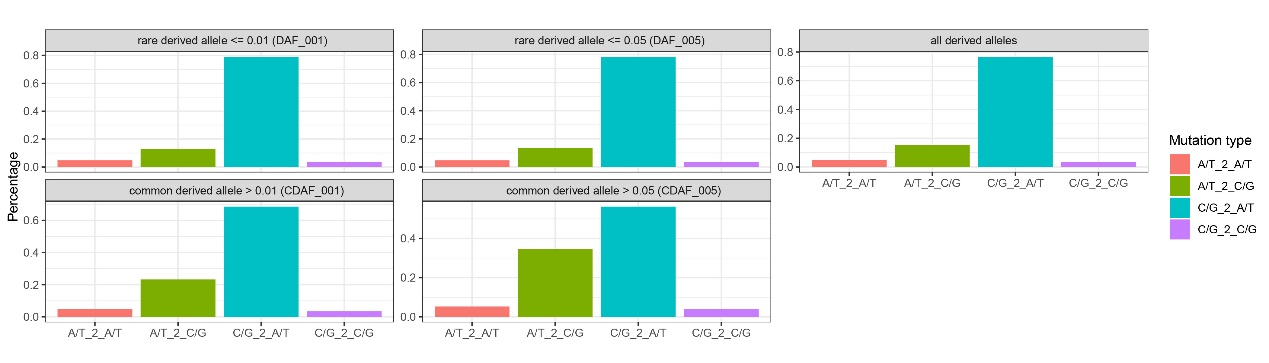

**Supplementary Figure 7.** Summary of mutation types in all derived alleles. Each panel represents loci within specific range of derived allele frequency. The plot shows high percentage of derived A/T alleles from ancestral C/G alleles (i.e. C/G_2_A/T), especially in loci with rare, derived alleles.

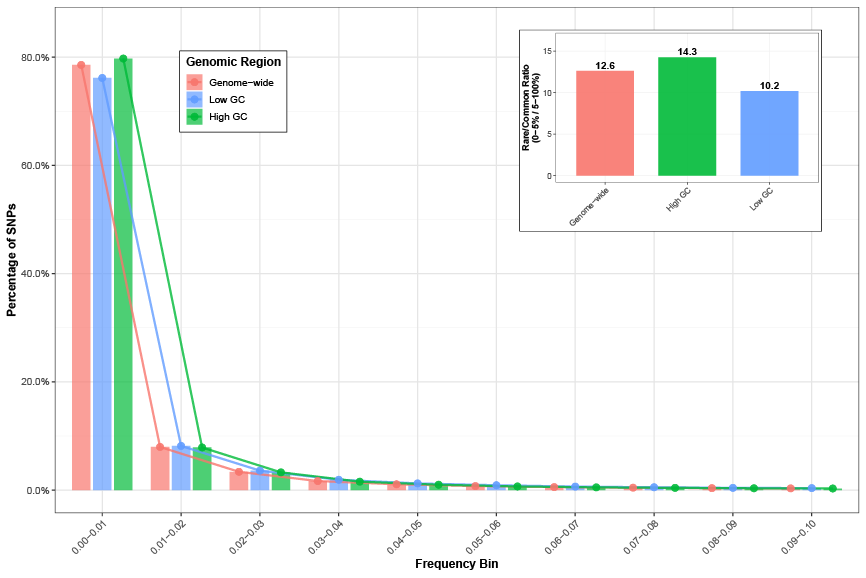

**Supplementary Figure 8.** Site frequency spectrum in different regions. Rare/common ratio of SNPs in the inset plot suggest more efficient selection in high GC regions than in low GC regions.

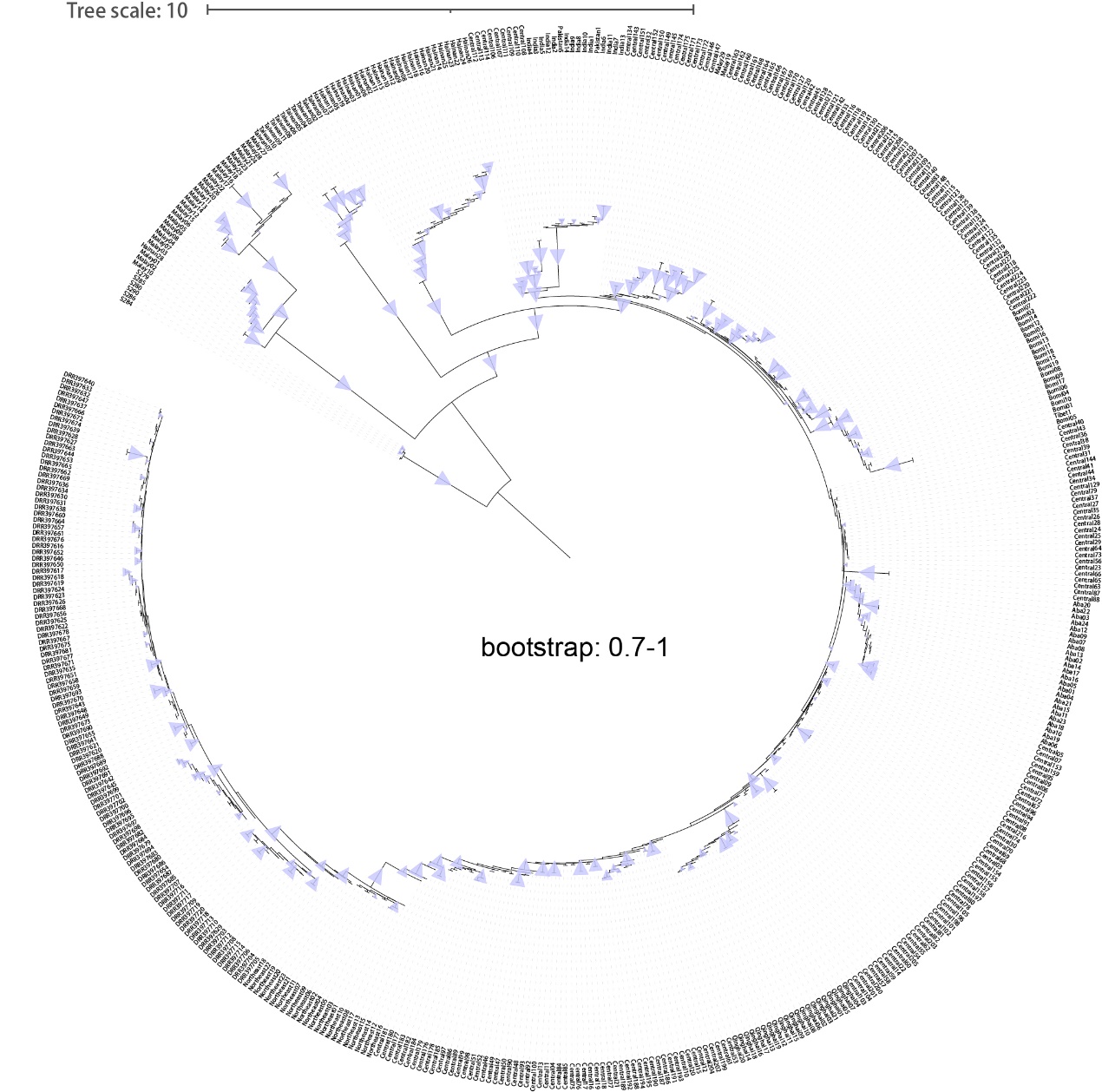

**Supplementary Figure 9.** Reconstruction of *Apis cerana* phylogeny based on nuclear polymorphisms in low GC regions. Bootstrap values over 0.7 were presented.

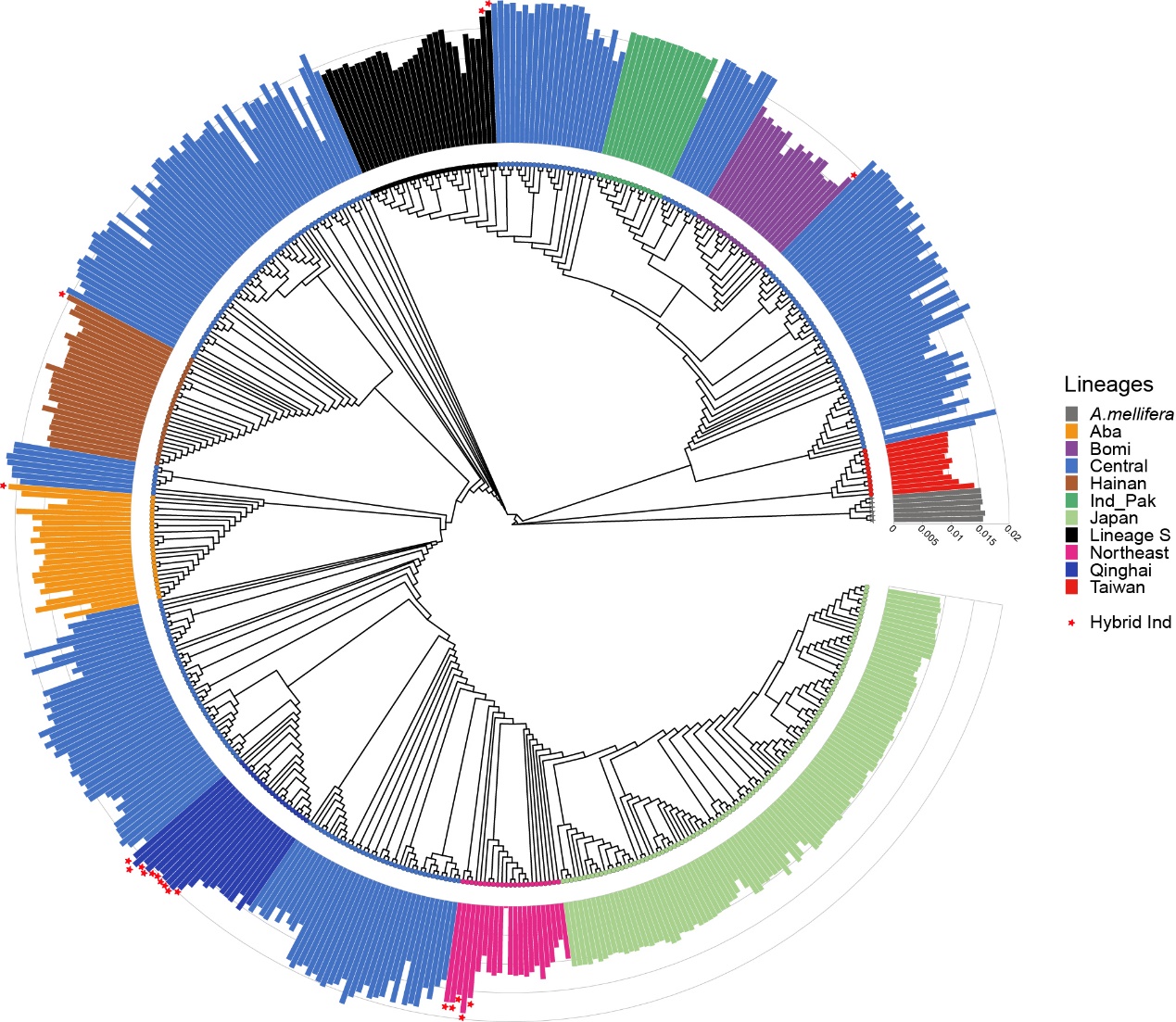

**Supplementary Figure 10.** Reconstruction of *Apis cerana* phylogeny based on nuclear polymorphisms in high GC regions. Het/Hom indicate ratio of homozygous vs. heterozygous sites in an individual. The hybrid individuals referred to genetic structure in Figure 1.
